# Mixing features of transcription factors and genes enables accurate prediction of gene regulation relationships for unknown transcription factors

**DOI:** 10.1101/2025.04.17.649264

**Authors:** Risa Okubo, Takashi Morikura, Yusuke Hiki, Yuta Tokuoka, Tetsuya J. Kobayashi, Takahiro G. Yamada, Akira Funahashi

**Affiliations:** Graduate School of Fundamental Science and Technology, Keio University, Kanagawa, Japan; Department of Biosciences and Informatics, Keio University, Kanagawa, Japan; Institute of Industrial Science, The University of Tokyo, Tokyo, Japan

## Abstract

Identifying regulatory relationships between transcription factors (TFs) and genes is essential to understand diverse biological phenomena related to gene expression. Recently, deep learning–based models to predict TFs that bind to genes from nucleotide sequences of the target genes have been developed, yet these models are trained to predict known TFs only. In this study we developed a deep learning model, GReNIMJA (Gene Regulatory Network Inference by Mixing and Jointing features of Amino acid and nucleotide sequences), to predict gene regulation even by unknown TFs. Our model is designed to mix the features of the TF amino acid sequences and nucleotide sequences of the target genes using a 2D LSTM architecture and to perform binary classification with the aim of determining the presence or absence of a regulatory relationship. The accuracy of our model in predicting regulatory relationships was 84.4% for known TFs (higher than those of conventional models) and 68.5% for unknown TFs; the latter is an unsolved task for conventional deep learning–based models. We expect our model to advance identification of unknown gene regulatory networks and contribute to the understanding of diverse biological phenomena.

## Introduction

Living organisms sustain their lives through the central dogma, where DNA is transcribed into mRNA, followed by the synthesis of proteins with amino acid sequences corresponding to the mRNA. During transcription, a transcription factor (TF) specifically binds to a transcription regulatory region and regulates gene expression near this region [1]. The complex relationships between TFs and genes form a gene regulatory network (GRN). Such networks are crucial in various biological phenomena, such as development [2, 3], stress responses [4, 5], and cancers [6, 7]. GRNs serve as blueprints for sustaining the biological phenomena, and therefore identifying GRNs is vital for understanding the various biological phenomena.

To identify GRNs, an experimental method called ChIP-seq has been widely used across various species and cell types [8–11]. ChIP-seq comprehensively detects the nucleotide sequences of genes where TFs bind, but it requires specific antibodies for each TF, making it labor-intensive and time-consuming as the TF number increases. To reduce experimental costs, statistical methods that use sequence motifs were developed [12]. The sequence motifs represent the occurrence probability of nucleotide sequences that are bound by specific TFs. Various sequence motifs have been registered in public databases such as JASPAR [13] and TRANSFAC [14]. Using these databases, statistical methods enable us to estimate the gene regulatory relationships for well-studied species like human and mouse. However, sequence motifs as such are insufficient for the accurate prediction of gene regulatory relationships because other factors, such as GC content and DNA shape, influence these relationships [15, 16]. In addition, these methods are fundamentally unable to predict gene regulatory relationships for unknown sequence motifs that have not been detected experimentally.

Deep learning–based discriminative models directly classify specific TFs from nucleotide sequences of the target genes [17, 18]. These models extract features from a nucleotide sequence with consideration of factors such as GC content and DNA shape, and predict the TFs that bind to that sequence [17, 18]. These models can predict the TFs that bind to genes even when the sequence motif they recognize is unknown. However, these models are trained to predict the specific TFs from genes, and thereby they are unable to predict gene regulatory relationships with unknown TFs. For identifying the GRNs composed of diverse TFs and genes, a deep learning-based model that can predict gene regulatory relationships even with unknown TFs is strongly required.

In this study we developed a deep learning–based model, GReNIMJA (Gene Regulatory Network Inference by Mixing and Jointing features of Amino acid and nucleotide sequences). A key point of this study is that GReNIMJA was designed, not to predict the specific TFs from genes, to predict whether the regulatory relationships exist or not from both of the amino acid sequences of TFs and nucleotide sequences of genes. Our model extracted features from both the amino acid sequences of TFs and the nucleotide sequences of target genes, mixed these features using a 2D LSTM architecture, and performed binary classification to predict the presence or absence of regulatory relationships. The accurate prediction of regulatory relationships even with unknown TFs by our model extends the frontier of the identification of unknown GRNs and may contribute to a deeper understanding of diverse life phenomena.

## Methods

### Dataset

To construct a dataset of TFs and their target genes, we collected the experimental data stored in the public DoRothEA [19] and Harmonizome database [20]. DoRothEA stores human and mouse gene regulatory relationships that were detected experimentally and statistically; it also groups these relationships by confidence level [19]. Harmonizome stores human gene regulatory relationships that were detected experimentally [20]. We collected the gene regulation data and obtained 2,674,639 pairs of human TFs and their target genes. We then obtained the TF amino acid sequences and their target gene sequences (upstream 1000 bp of each gene) from the public genome database GenBank [21] (accession number for the human genome: GCF_000001405.39) and constructed pairs of TF amino acid sequences and gene nucleotide sequences. We removed pairs that were duplicated in the DoRothEA and Harmonizome databases, contained unidentified residues or bases, or were assigned to the lowest confidence level in DoRothEA, and obtained 1,806,075 pairs (301,915 from DoRothEA and 1,504,160 from Harmonizome). The constructed dataset contained 697 TFs. We assigned positive labels to the pairs for which the gene regulatory relationship was indicated as presence in the database (Supplementary Figure S1), then shuffled the TFs and genes and assigned negative labels to the shuffled pairs for which the gene regulatory relationship was not detected. The same number of nucleotide sequences as the number of genes regulated by TFs was randomly selected from genes other than the regulated genes.

Our gene regulation dataset (3,383,776 pairs) consisted of 1,806,075 positive pairs and 1,577,701 negative pairs; 316,298 pairs were excluded and used as test data. The remaining dataset was divided into training data and validation data (5-fold cross validation). At each fold, the model with the lowest value of the loss function at each epoch was selected, followed by selection of the best model in the whole dataset, and its performance was evaluated using the test data.

### Embedding of amino acid sequences and nucleotide sequences

To convert amino acid sequences and nucleotide sequences into numerical vector representations, each sequence was converted into a *k*-mer representation (Supplementary Figure S2a), which was fed to each specifically pre-trained word2vec model [22] for human amino acid sequence and nucleotide sequences(Supplementary Figure S2b). The sequence length *k* was 4 for amino acid sequences [23] and 5 for nucleotide sequences [24].

Word2vec learns mapping from strings to numerical vectors while considering the semantic information of words within sentences [22]. Word2vec is useful for predicting biological semantics such as protein family attributes and protein–protein interactions [25–27]. To predict surrounding words from the target word, we pre-trained word2vec with the skip-gram algorithm. The word2vec architecture is constructed as a neural network with the input, middle, and output layers that are fully connected. The softmax function is applied to the output layer, and the parameters of the model are updated by calculating the total cross-entropy loss between the predicted vector and the ground-truth vector of surrounding words. The number of dimensions *d* of the numerical vectors in the output layer was set to 100 for amino acid sequences [23] and 50 for nucleotide sequences [24]. The number of epochs for training the word2vec model was set to 10, and the maximum distance between the predicted surrounding words and the target word was set to 10 words. We trained two word2vec models, one with amino acid sequences and the other with nucleotide sequences from GenBank [21] (accession number: GCF_000001405.39). These models do not learn the presence or absence of gene regulatory relationships, but just functions as a pre-trained model that performs embedding, so we trained the embedding model on all of the above sequence data registered in GenBank.

To obtain a numerical matrix, we repeatedly input the sub-sequences of the *k*-mer representation into each pre-trained word2vec model, and combined the numerical vectors from each *k*-mer representation (Supplementary Figure S2c).

### Overview of the proposed model

To accurately predict gene regulatory relationships even with unknown TFs, we developed a deep learning–based model, GReNIMJA (Gene Regulatory Network Inference by Mixing and Jointing features of Amino acid and nucleotide sequences), which: (i) extracted each feature from numerical matrices with the amino acid sequences and nucleotide sequences embedded using pre-trained word2vec; and (ii) mixed these features using a 2D LSTM architecture (Figure 1). It used binary classification to predict the presence or absence of regulatory relationships.

**Figure 1:**
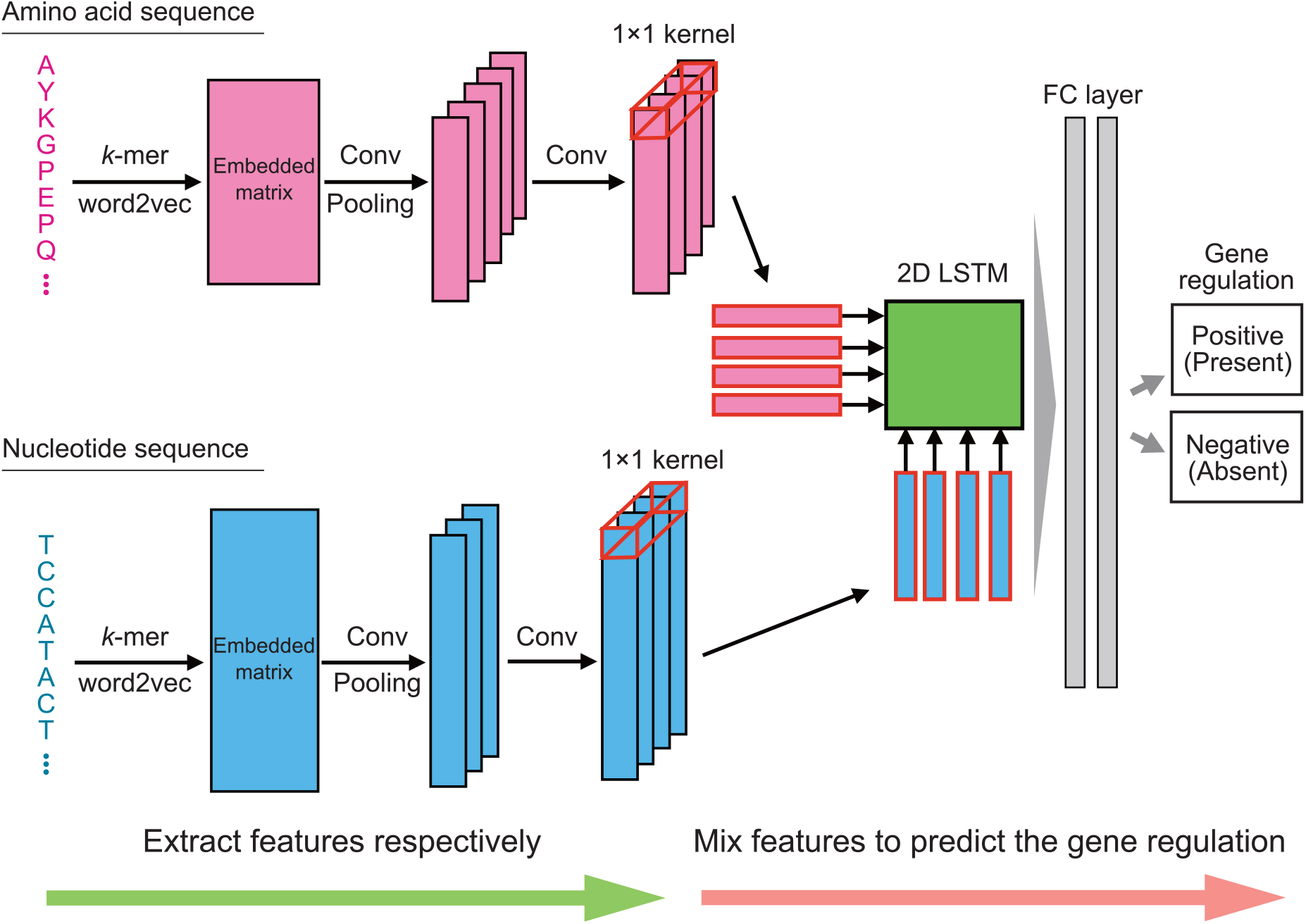
Overview of the architecture of the proposed model The amino acid sequence of a transcription factor (TF) and the nucleotide sequence of its target gene are converted into *k*-mer representations. Pre-trained word2vec models are used to embed these representations into numerical matrices. The convolution (Conv) and pooling layers are used to extract the features of this matrix. To align the dimensions of the extracted features, the features are input to the convolution layer with a 1 *×* 1 kernel. The aligned features are mixed in 2D LSTM. The output from the 2D LSTM is input to the fully connected (FC) layers with sigmoid function. Using the feature output from the FC layers, our model classifies the regulatory relationship between the TF and gene as positive (present) or negative (absent).

(i) Each of the embedded representations was input to the convolution layer and pooling layer with a dropout process that randomly set the parameter values to zero. Because the dimensions of the features extracted from the amino acid sequences and nucleotide sequences were different, we performed convolution processing using a 1 × 1 filter for each feature to align the number of dimensions. (ii) The two-dimensional Long Short-Term Memory (2D LSTM) model enables mixing different series of data; it was used to mix both features [28]. The recursive memory unit structure of LSTM aimed to memorize long sequences by updating internal states; it had an input gate, an output gate, a memory cell to store the state, and an oblivion gate to determine whether to use the features stored in the memory cell. Although LSTM is able to store sequences for long periods, it handles only one kind of series as input. We incorporated 2D LSTM into the proposed model to handle two series as inputs: the TF amino acid sequence and the gene nucleotide sequences (Supplementary Figure S3). To determine how to mix the two series of sequences, we added a lambda gate to the conventional LSTM and obtained the memory unit structure of 2D LSTM. The features mixed by the 2D LSTM were input into the two fully connected layers with sigmoid function to predict the presence or absence of the gene regulatory relationship.

### Training procedure

The number of epochs was fixed at 100. Mini-batch size was set to 1024 for the training dataset and 512 for the validation dataset. As the loss function, a binary cross-entropy loss was used. Adam [29] was used as the optimizer; the initial learning rate for Adam was set to *lr* = 0.00001. Hyperparameters of GReNIMJA were tuned using Optuna [30] that implemented Bayesian optimization with the Tree-structured Parzen Estimator algorithm (Supplementary Table S1). For model training, we used NVIDIA V100 32GB PCIe (FP32 14 TFLOPS) and NVIDIA A100 40GB PCIe (FP32 19.5 TFLOPS).

### Evaluation of model performance for known transcription factors

Using the test dataset, we evaluated the prediction with the Accuracy, AUROC, AUPR, F-measure, and Matthews Correlation Coefficient (MCC) metrics as follows:

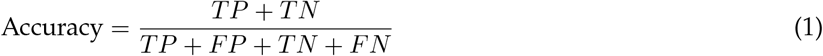

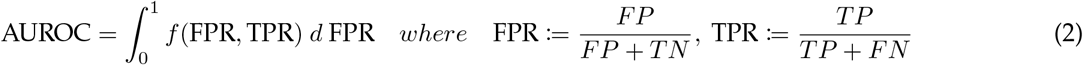

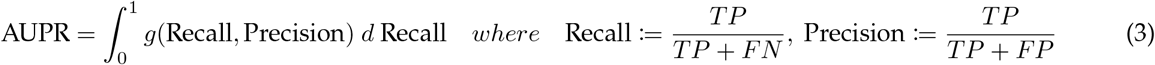

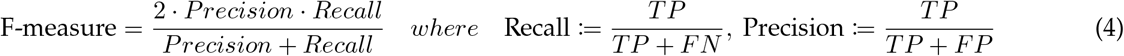

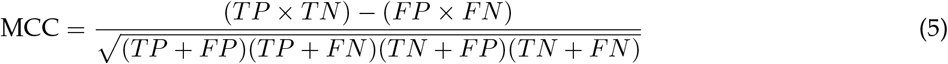

where *TP*, true positive; *TN*, true negative; *FP*, false positive; *FN*, false negative; FPR, *FP* rate; and TPR, *TP* rate. Precision and Recall evaluate the number of false positives and false negatives, respectively. AUROC represents the area of the ROC curve *f* (FPR, TPR) and evaluates the robustness of the model by adjusting the balance between FPR and TPR. The function *f* represents the curve connecting the sample points of FPR and TPR. AUPR represents the area of the PR curve *g*(Recall, Precision) and evaluates the robustness of the model by adjusting the balance between Precision and Recall. The function *g* represents the curve connecting the sample points of Precision and Recall. The F-measure evaluates the balance between false positives and false negatives. MCC evaluates the accuracy of a model in a binary classification task by adjusting the balance between positive and negative cases when the number of one class is too small. When all predictions are wrong, MCC is − 1; when predictions are made at random, MCC is 0; and when all predictions are correct, MCC is 1. We calculated the average of the evaluation metrics for all test data as a representative value. For Accuracy, AUROC, and AUPR [31], we also calculated the average of the evaluation metrics for each TF in the test data.

We compared the performance of GReNIMJA with that of a traditional statistical analysis method (PWMScan [12]), classical machine learning methods (Support Vector Machine (SVM) [32], Logistic Regression (LR) [33], and eXtreme Gradient Boosting (XGBoost) [34]), a conventional deep learning method (DeepRAM [31], and a multi-class classification model TBiNet [35]) on the same dataset.

PWMScan predicts TF binding from motifs in the target nucleotide sequences. The motifs were obtained from the JASPAR database [13]; it stores position frequency matrices (PFMs) that represent the probability of nucleotide sequence binding to a known TF. The PFM was converted into a position probability matrix (PPM; 6). The PPM was converted into a position weight matrix (PWM) that corrected the bias in the occurrence probability of nucleotide sequences across the whole genome(Supplementary Figure S4).

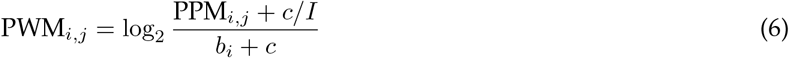

where PPM_*i,j*_ and PWM_*i,j*_ represent the occurrence probability of PPM and PWM at each position *i, j*. Parameter *b*_*i*_ represents the occurrence probability of the nucleotide sequence in the whole genome. Parameter *c* s a constant value introduced for computational stability, and was set to 1 [36]. Parameter *I* represents the sequence length, and was set to 4 for nucleotide sequences and 20 for amino acid sequences. For PWMScan performance evaluation, TFs not included in the JASPAR database were excluded from the test dataset.

SVM learns a binary classification to maximize the distance between the decision boundary and each data point [32]. Generalized linear model LR follows a Bernoulli distribution and learns a binary classification based on the probability that a data point belongs to a specific class [33]. XGBoost (a gradient boosting decision tree model) is an ensemble model of decision trees that learns a binary classification [34]. In these models, the length of the input numerical vector must be fixed. In accordance with [37], we fixed the length of input numerical vector by obtaining the weighted average of each numerical vector that was converted from *k*-mer representation by using the pre-trained word2vec model. Values of term frequency-inverse document frequency consider the importance of each *k*-mer [37] and were used as weights for averaging.

DeepRAM [31] classifies whether a specific TF is a regulatory factor or not for the input nucleotide sequences. DeepRAM architecture consists of a convolution layer and LSTM. Because DeepRAM learns a binary classification task, training multiple models for each TF is required; because of computational constraints, we randomly selected 44 TFs of the 697 TFs in the dataset, and trained each model for each TF. TBiNet [35] predicts the presence or absence of the gene regulatory relationship for each TF from the input nucleotide sequences. TBiNet architecture consists of convolution layers, LSTM, and an attention layer. Because TBiNet learns a multi-class classification task, TBiNet cannot predict TFs outside of the training dataset. We trained TBiNet on TFs from the test dataset. The hyperparameters for each TF in each DeepRAM model were optimized by using Bayesian optimization with the Tree-structured Parzen Estimator algorithm described in [31]. We used the hyperparameters for TBiNet described in [35].

### Evaluation of model performance for unknown transcription factors

We divided 697 TFs as follows: training, 597; validation, 30; test, 70. We trained the model with 50 epochs and used the Accuracy, AUROC, and AUPR metrics to evaluate its performance. We also compared the performance of our model with that of classical machine learning methods using the same datasets. A traditional statistical analysis method and deep learning methods were excluded from the comparison because they cannot predict regulatory relationships for unknown TFs.

### Evaluation of model robustness

The performance of deep learning models generally depends on the number of datapoints; when this number is small, the performance is often degraded [38, 39]. To evaluate the robustness of our model for the number of datapoints for TFs, we analyzed the accuracy of prediction for that number.

The performance of LSTM can also be degraded for extremely long sequences [40]. Although most TFs in the dataset had around 500 amino acids, the longest one had 4005 (Supplementary Figure S5). We analyzed the accuracy of prediction for each sequence length. To quantify the effect of the length of the TF amino acid sequence on prediction bias, we analyzed the true positive, true negative, false positive, and false negative predictions by our model for each sequence length.

### Feature analysis of the trained model

To interpret GReNIMJA, we analyzed features in (i) the convolution layer and (ii) the last output layer of the model. (i) We analyzed the activated amino acid sequences and nucleotide sequences in our model. The analysis algorithm was based on that used in DeepBind [17]. Briefly, all sequences in the training dataset were input into the convolution layer, and the feature vectors *u* were obtained. By adding all features and subtracting the mean value of the feature vector u_*mean*_, we obtained the centered feature vector *v* := *u* − u_*mean*_. The reference value Z was calculated as follows:

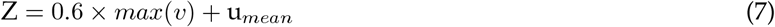

An array corresponding to the position *>* Z was extracted as the active array. PWM was calculated for the active array to obtain the weighted occurrence probability of the sequence elements. To analyze amino acid sequence activation, we searched for the most similar motif by comparing the calculated PWM with the PWM stored in the PROSITE database [41]. For motif searching, we used the Tomtom database [42]. To analyze nucleotide sequence activation, we calculated the total sum of the corrected occurrence probabilities for each nucleotide in the PWM of each filter and chose the top two nucleotides with the highest total values. The number of the obtained nucleotides was counted, and the occurrence ratio of the nucleotides was calculated. (ii) We reduced 140 dimensions of the features output by the model using the test dataset to 2 dimensions using t-Distributed Stochastic Neighbor Embedding (t-SNE) [43]. By plotting the reduced features in a 2-dimensional space, we qualitatively evaluated the features of the decision boundary regarding the presence or absence of gene regulatory relationships.

To evaluate whether the mixture of features contributed to the prediction performance, the decision boundaries for the presence or absence of the gene regulatory relationship were visualized (Supplementary Figure S6a). To improve visibility, we visualized each plot in the dimensionality-reduced space for 10 TFs and 100 target genes selected randomly from the test dataset. If the features were not mixed by the model, all dots should be concentrated at the same point without crossing the decision boundary (Supplementary Figure S6b). If the features were mixed, all dots should be distributed across the decision boundary (Supplementary Figure S6c).

Our model attempts to predict gene regulatory relationships for unknown TFs by using feature similarity to the learned sequence. To evaluate whether the TF features learned by the model reflect the similarity between TFs, we analyzed sequence similarity between a TF and its neighboring TFs on the plot in the dimensionality-reduced space using Basic Local Alignment Search Tool (BLAST) [44]. To simplify the analysis, we selected the TFs with a prediction accuracy of ≥20% higher by our model than those of conventional deep learning models. We visualized the relationship between the number of datapoints and the proportion of neighboring TFs homologous to the TF of interest. We also analyzed sequence similarity between the unknown TFs present only in the test dataset and their neighboring TFs, and calculated the percentage of TFs in the neighborhood homologous to the unknown TF.

## Results

### Proposed model outperformed conventional models in predicting gene regulatory relationships for known transcription factors

The proposed model GReNIMJA outperformed conventional models in all evaluation metrics (Table 1). Its highest accuracy (0.844) indicated that the model was the most accurate at predicting gene regulatory relationships. The highest F-measure (0.847) indicated the best balance between false negatives and false positives. The highest AUROC (0.928) and AUPR (0.929) indicated that the model was more robust than the conventional models. The MCC value of GReNIMJA was also the highest (0.689); MCC evaluates precision while adjusting the balance of the number of datapoints. Because the number of datapoints in the constructed dataset was larger for positive data than for negative data, GReNIMJA had high precision in predicting not only positive but also negative data. In summary, GReNIMJA outperformed conventional models in terms of prediction accuracy, balance between false negatives and false positives, robustness, and precision.

**Table 1:**
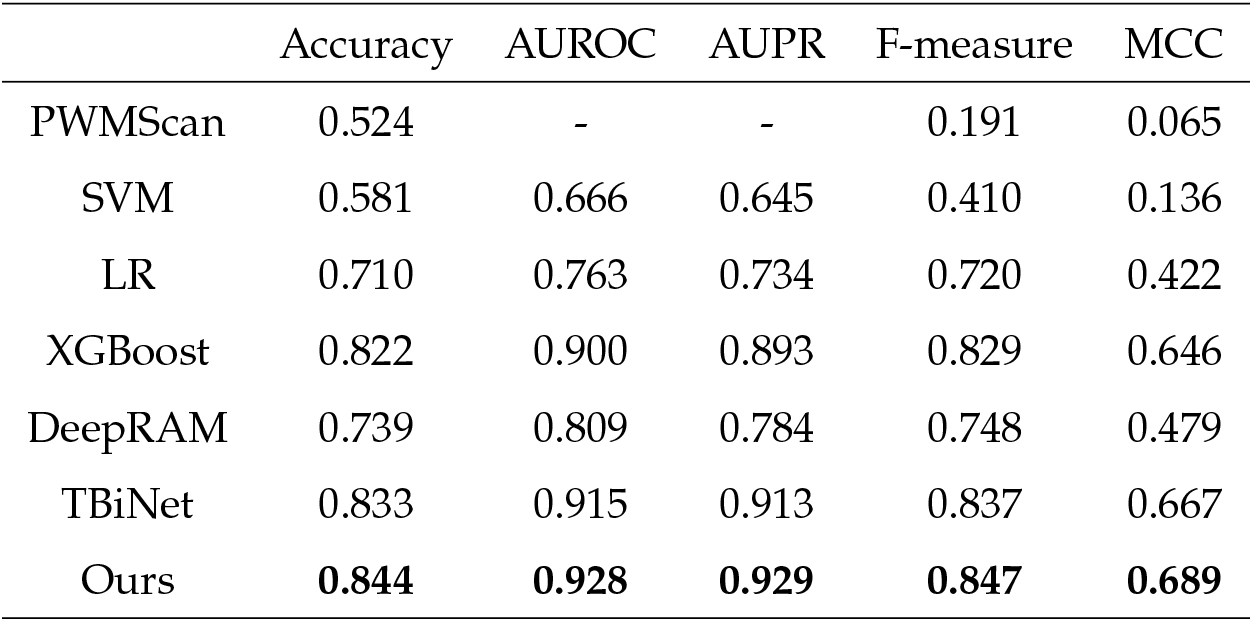
Evaluation of model performance in the prediction of gene regulatory relationships for known TFs Each row shows values of evaluation metrics for the indicated model. The average values of each metric for the test data are shown. The highest values are in bold. PWMScan does not allow calculation of AUROC and AUPR.

The number of datapoints for TFs tended to decrease as the sequence length increased (Supplementary Figure S5). To evaluate model performance for TFs of different length, we analyzed the accuracy, AUROC, and AUPR for each TF. GReNIMJA outperformed the conventional models in all evaluation metrics for each TF (Table 2). Accuracy, AUROC, and AUPR of the prediction by GReNIMJA for all TFs were 0.011, 0.013, and 0.016 points higher, respectively, than those of TBiNet, which showed the highest performance among the conventional models (Table 1). For each TF, these metrics were 0.093, 0.058, and 0.049 points higher in GReNIMJA than in TBiNet (Table 2). Thus, the degree of performance improvement relative to the conventional models was higher for each TF than for all TFs. These findings and the distribution of the sequence lengths (Supplementary Figure S5) suggested that GReNIMJA predicts the gene regulatory relationships more accurately and robustly even for the TFs with few datapoints and long sequences. Thus, GReNIMJA would be useful for the identification of regulatory relationships for various TFs and genes.

**Table 2:** Evaluation of model performance for each known TF in the prediction of regulatory relationships Each row shows values of evaluation metrics for the indicated model. A representative value was calculated as the average metric value for each TF; the average representative values were then calculated. The highest values are in bold. PWMScan does not allow calculation of AUROC and AUPR.

|  | Accuracy | AUROC | AUPR |
| --- | --- | --- | --- |
| PWMScan | 0.521 | - | - |
| SVM | 0.544 | 0.644 | 0.662 |
| LR | 0.612 | 0.669 | 0.677 |
| XGBoost | 0.674 | 0.766 | 0.764 |
| DeepRAM | 0.639 | 0.690 | 0.663 |
| TBiNet | 0.658 | 0.782 | 0.790 |
| Ours | **0.751** | **0.840** | **0.839** |

### Proposed model was robust against the small number of datapoints and long sequences of transcription factors

The relationship between the number of datapoints and the prediction accuracy of our model is shown in Figure 2. Prediction accuracy tended to deteriorate with the decrease in the number of datapoints, but remained higher than that of conventional models at a number of datapoints <1000 (Figure 2a). In addition, the accuracy of our model was not dependent on sequence length (Figure 2b). The proportion of positive (true or false) and negative (true or false) predictions was similar regardless of sequence length (Figure 2c). This suggests that sequence length do not affect the bias of our model. Overall, GReNIMJA was able to robustly predict gene regulatory relationships even for the TFs with a small number of datapoints and long sequences.

**Figure 2:**
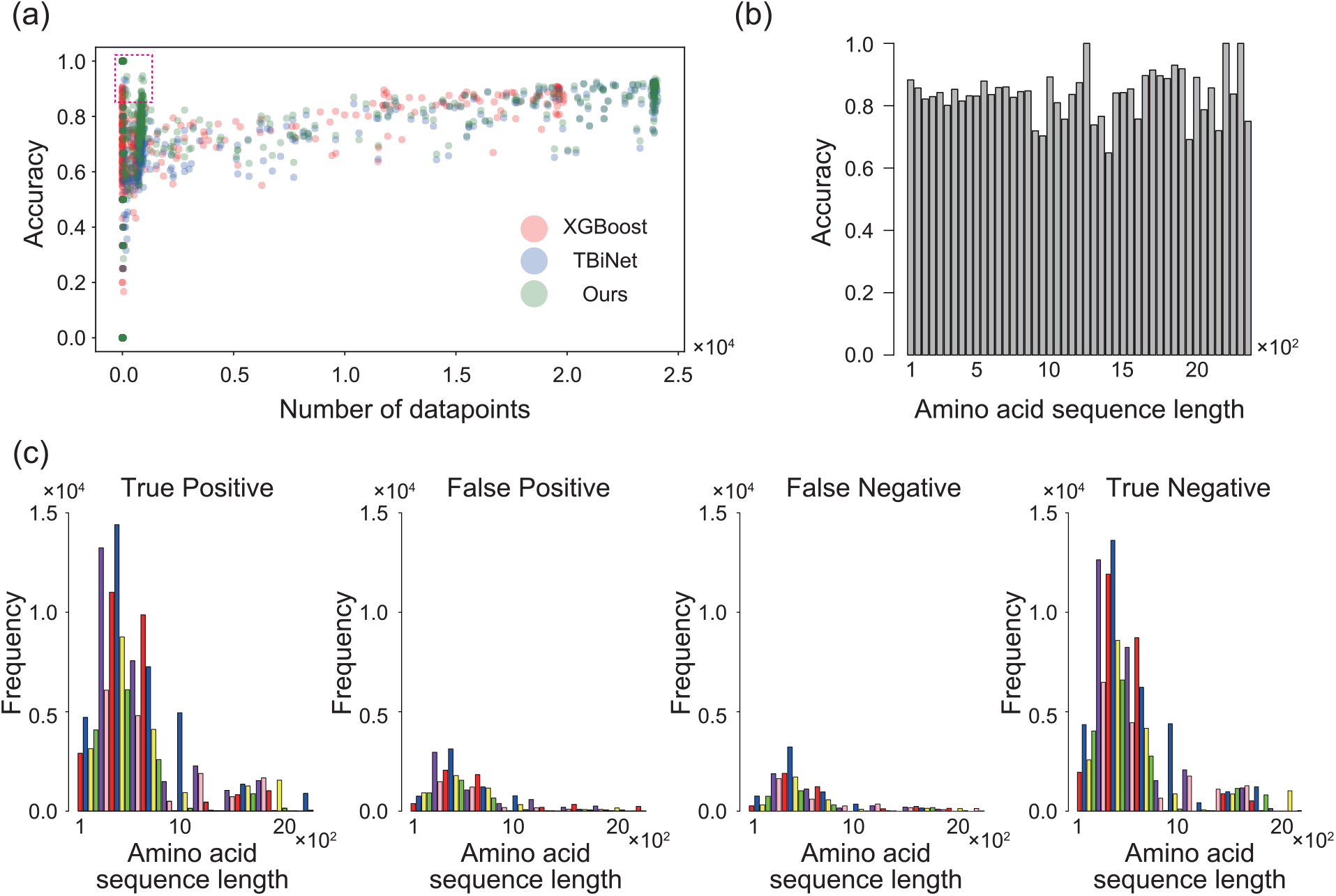
Effect of the number of datapoints and TF length on model performance (a) The horizontal axis represents the number of TFs in the training dataset, and the vertical axis represents the accuracy for each TF in the test dataset. The area surrounded by the dotted line represents the region where the performance of our model exceeded that of the conventional model for a TF with <1000 datapoints. (b) Average accuracy according to TF sequence length. The groups of sequence lengths were constructed in increments of 50 residues. (c) The relationship between TF length and TF frequency predicted in the indicated categories. The groups of sequence lengths were constructed in increments of 50 residues.

### Proposed model acquired biological context related to gene regulatory relationships

To interpret the basis of the predictions by our model, we analyzed the features of the convolution and last output layers. In the convolution layer, we found that a total of 11 types of TF domains were activated, including leucine zippers and zinc fingers (Table 3) In nucleotide sequences, our model focused on AT content (Table 4). In the last output layers, a dimensionality reduction analysis by t-SNE showed that the positive and negative data were concentrated in the same region and formed clusters (Supplementary Figure S7). Our model might improve performance as a result of learning a decision boundary that forms easily separable clusters. It is noteworthy that it learned the decision boundary of a gene regulatory relationship by using the biological domain context as a basis for the prediction, even though the model was not explicitly given any biological knowledge that was important for gene regulatory relationships.

**Table 3:**
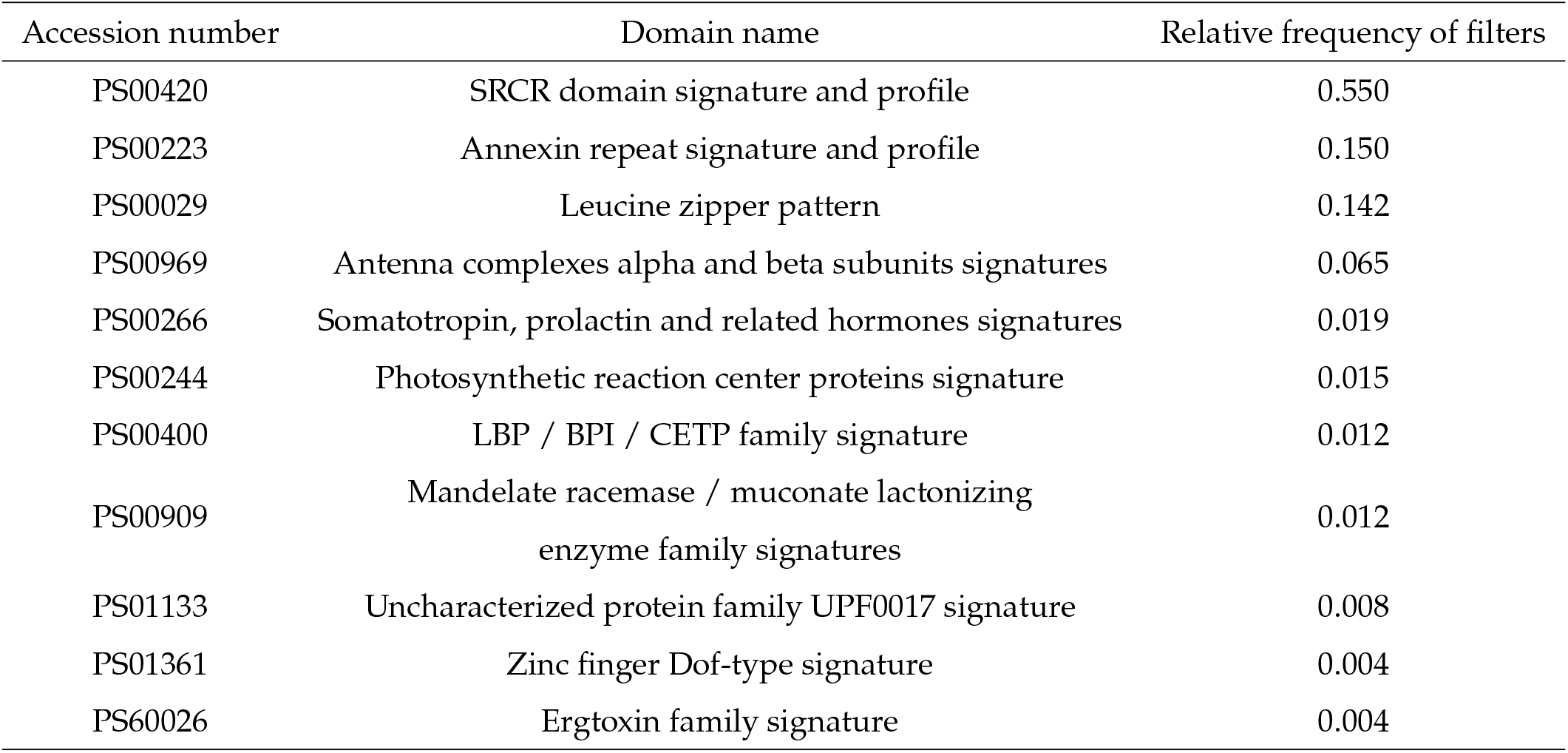
Activated amino acid sequences in the convolution layer. The relative frequency of each sequence in the convolution filter is shown in the right-most column. Detailed information on the approach is provided in the Methods section.

| Accession number | Domain name | Relative frequency of filters |
| --- | --- | --- |
| PS00420 | SRCR domain signature and profile | 0.550 |
| PS00223 | Annexin repeat signature and profile | 0.150 |
| PS00029 | Leucine zipper pattern | 0.142 |
| PS00969 | Antenna complexes alpha and beta subunits signatures | 0.065 |
| PS00266 | Somatotropin, prolactin and related hormones signatures | 0.019 |
| PS00244 | Photosynthetic reaction center proteins signature | 0.015 |
| PS00400 | LBP / BPI / CETP family signature | 0.012 |
| PS00909 | Mandelate racemase / muconate lactonizing<br>enzyme family signatures | 0.012 |
| PS01133 | Uncharacterized protein family UPF0017 signature | 0.008 |
| PS01361 | Zinc finger Dof-type signature | 0.004 |
| PS60026 | Ergtoxin family signature | 0.004 |

**Table 4:** Relative frequencies of activated nucleotide pairs in the convolution layer. The highest value is in bold. Detailed information on the approach is provided in the Methods section.

| AC | AG | AT | CG | CT | GT |
| --- | --- | --- | --- | --- | --- |
| 0.019 | 0.004 | **0.958** | 0.015 | 0.004 | 0.000 |

To evaluate the effect of feature mixing on model performance, we visualized the relationship between the decision boundary and feature plots randomly selected from the test dataset in the dimensionality-reduced space. If the features were not mixed, the embedded numerical vectors of amino acid sequences and nucleotide sequences were aggregated in the same region, but if the features were mixed, each feature of TF or gene was distributed across the decision boundary. The feature dots of the TFs tended to form clusters across the decision boundary (Figure 3a), whereas the feature dots of the genes were sparsely scattered across the decision boundary (Figure 3b). This suggests that our model considers the interaction between TFs and genes when predicting the regulatory relationships between them, in agreement with biological knowledge.

**Figure 3:**
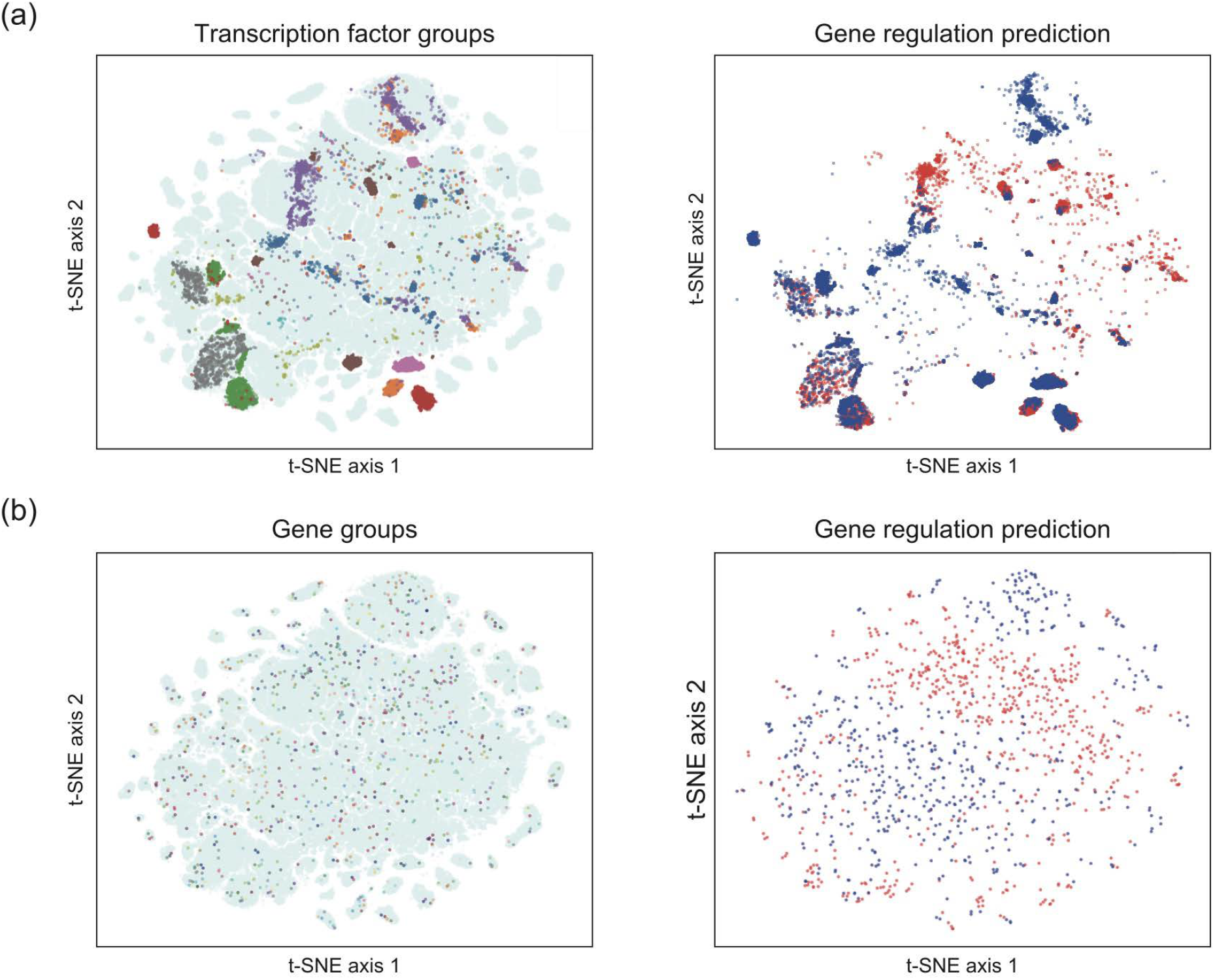
Effect of feature mixing on the prediction of gene regulatory relationships by our model Dimension reduction of the features obtained from the output layer using t-SNE was performed. Each dot represents features corresponding to each TF and gene. Left panels, randomly selected (a) TFs or (b) genes. Dots of different colors represent different TFs or genes; light blue, TFs or genes that were not selected. Right, prediction results for the gene regulatory relationships for each feature. Blue, features predicted as positive; red, features predicted as negative. Detailed information about the approaches used is provided in the Methods section.

We designed our model to predict gene regulatory relationships by using feature similarity to the learned sequence even if the number of datapoints for a TF is small and this TF has not been learned. To evaluate whether the learned features of a TF preserve the TF characteristics, we analyzed sequence similarity between a TF and its neighboring TFs on the plot in the dimensionality-reduced space. For TFs with few datapoints, neighboring TFs had high similarity to the TF of interest (Figure 4a). For unknown TFs, neighboring TFs had high similarity at a rate of 10%–20% for the TF of interest (Figure 4b). The ability of our model to make accurate predictions even for TFs with a small number of datapoints and for unknown TFs ((Figure 2, Table 5) indicates that its high accuracy was achieved by using features similar to the learned TFs.

**Table 5:** Evaluation of model performance for each unknown TF in the prediction of regulatory relationships Each row shows evaluation metric values for the indicated model. A representative value was calculated as the average value for each TF; the average representative values are shown. The highest values are in bold. Detailed information on the evaluation metrics and comparison is provided in the Methods section.

|  | Accuracy | AUROC | AUPR |
| --- | --- | --- | --- |
| SVM | 0.548 | 0.600 | 0.616 |
| LR | 0.559 | 0.620 | 0.626 |
| XGBoost | 0.651 | 0.683 | 0.687 |
| Ours | **0.685** | **0.698** | **0.694** |

**Figure 4:**
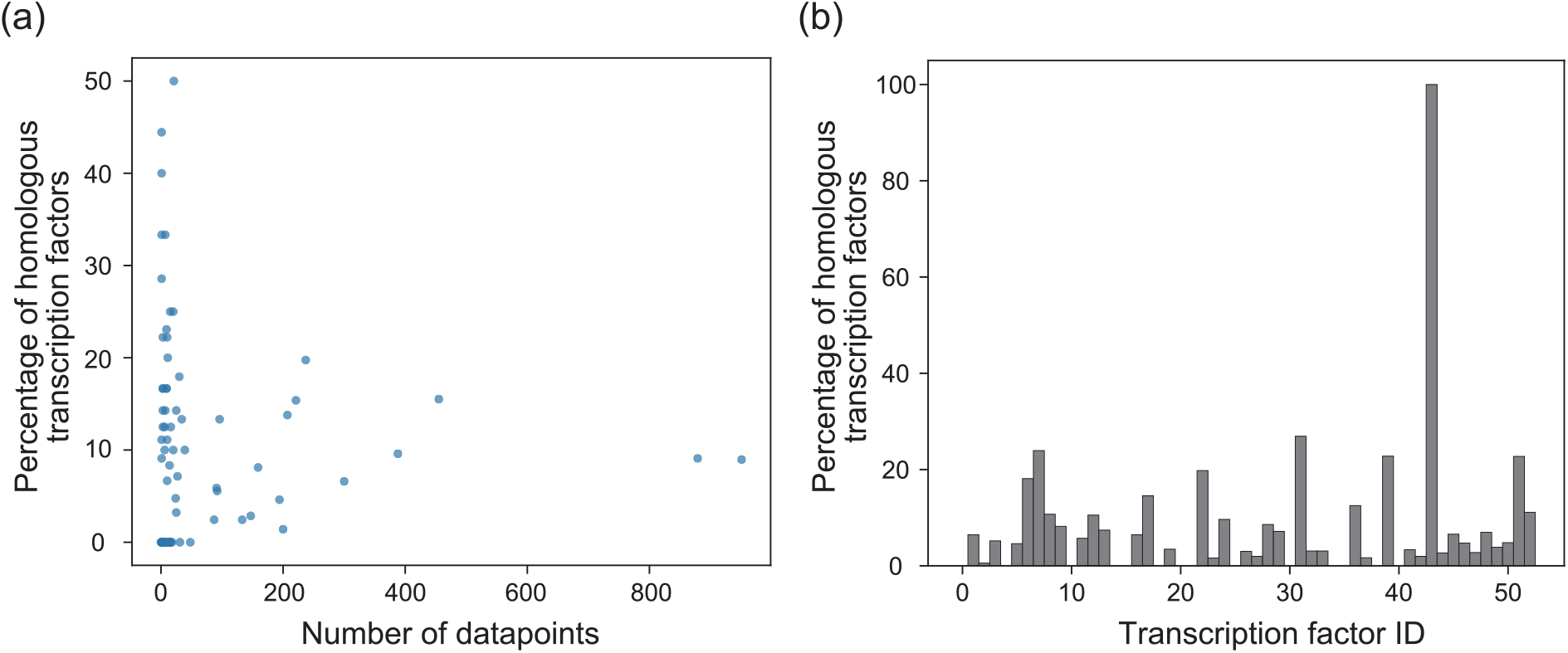
Homology analysis. (a) Relationships between the number of datapoints for the TF and the percentage of neighboring TFs homologous to the TF of interest in the training dataset. The TFs of interest were selected from among those with *>*1000 datapoints and those with the gene regulatory relationships predicted with an accuracy of ≥20% higher than that of TBiNet. Each dot represents one TF. (b) Proportion of neighboring TFs found to be homologous to the unknown TF of interest. The IDs of unknown TFs were assigned randomly.

Overall, our model achieved highly accurate predictions of gene regulatory relationships by considering the interactions between TFs and genes, focusing on the DNA-binding domains of the TFs and AT content of the genes, and using features similar to the learned TFs.

### Our model achieved high-precision predictions even for unknown transcription factors

Our model accurately predicted gene regulatory relationships for unknown TFs (Table 5). In all evaluation metrics, its performance was higher than that of the traditional statistical analysis methods and classical machine learning methods (Table 5). The highest accuracy of our model (0.685) indicated that it most accurately predicted gene regulatory relationships. The highest AUROC (0.698) and AUPR (0.694) of our model indicated that it can most robustly estimate gene regulation relationships. Because our model can infer gene regulatory relationships accurately and robustly for unknown TFs, it can be useful for the identification of diverse gene regulatory relationships, in particular for TFs that have not even been detected experimentally.

## Discussions

In this study, we developed a deep learning–based model, GReNIMJA, that is able to predict gene regulatory relationships for unknown TFs (Figure 1). GReNIMJA predicted gene regulatory relationships for known TFs better than did the conventional models (Table 1, 2). The model was robust to the number of datapoints and TF sequence length (Figure 2). By analyzing the features of GReNIMJA, we revealed that it predicted gene regulatory relationships by using mixed information on TFs and target genes (Figure 3). As a basis for the prediction, GReNIMJA learned the biological domain context, namely the DNA-binding domains in the TFs and the AT content of the target genes (Table 3, Table 4). Homology analysis revealed high similarity between TFs that had similar features learned by GReNIMJA (Figure 4). The model might have learned latent representations that reflect the homology among TFs. By leveraging the latent representations of similar pre-trained TFs, the model was able to predict gene regulatory relationships with high accuracy even for unknown TFs. Consistent with this result, we showed that GReNIMJA predicted the regulatory relationships of unknown TFs with high performance (Table 5). Prediction of gene regulatory relationships requires prediction of not only the binding between TFs and genes, but also whether the TFs regulate these genes or not [45]. Although conventional deep learning–based models are able to accurately predict whether or not a specific TF and gene will bind to each other [17, 18], GReNIMJA can also predict the presence or absence of gene regulatory relationships on the basis of biological domain knowledge. This fundamental difference might have contributed to the improved performance.

We found that PS00029 (leucine zipper) and PS01361 (zinc finger) were activated in the convolution layer (Table 3); both are DNA-binding domains [46]. The scavenger receptor cysteine-rich (SRCR) domain was also activated. This domain is present in cell-surface and secreted proteins mainly related to the immune system [47], and it mediates protein–protein interactions [48]. As the binding specificity of TFs can be changed through interactions with other proteins [49], the SRCR domain might affect gene regulatory relationships through such interactions. Together, these findings suggest that the performance of GReNIMJA was improved by its ability to recognize the DNA-binding domains, which are important in gene regulation, and the SRCR domain, which affects gene regulatory relationships through protein–protein interactions.

We found that ATs in nucleotide sequences were activated in the convolution layer (Table 4). Analysis of the nucleotide content at binding sites and non-binding sites of TFs has revealed that TFs have binding sequence preferences; some TF families tend to bind to AT-rich or GC-rich sequences [50]. These considerations suggest that the performance of GReNIMJA was improved by its ability to recognize whether the gene nucleotide sequence was AT-rich or not.

Predictions by GReNIMJA were based only on sequence information, but gene regulatory relationships are also influenced by the three-dimensional structure of DNA and geometric variations in TF structures [51], as well as hydrophobicity [52]. Because the physical characteristics that govern higher-order functions related to DNA and proteins represent fundamental mechanisms shared across organisms [53], the integration of such information might be useful to predict gene regulatory relationships more generally. By generalizing GReNIMJA, we may be able to obtain gene regulatory relationships as blueprints underlying biological phenomena, paving the way for a deeper understanding of diverse life processes.

In summary, we demonstrated that GReNIMJA predicts regulatory relationships between TFs and genes with high accuracy and robustness, even for unknown TFs. We expect this model to contribute to a deeper understanding of biological phenomena by accurately identifying complex gene regulatory relationships that have not yet been fully elucidated.

## Code availability

The source code developed in this study is available from https://github.com/funalab/GReNIMJA.

## Data availability

The data used in this paper is available at https://github.com/funalab/GReNIMJA.

## Acknowledgements

The research was funded by JST CREST, Japan Grant Number JPMJCR2011 to T.J.K. and A.F.

## Author contributions

R.O., Y.H., Y.T., T.G.Y. and A.F. designed the conceptual idea and the study. R.O. implemented the proposed algorithm. R.O., T.M., Y.H., Y.T., T.J.K., T.G.Y. and A.F. wrote the manuscript, with suggestions from the other authors.

