## Supplementary Material for "Mixing features of transcription factors and genes enables accurate prediction of gene regulation relationships for unknown transcription factors"

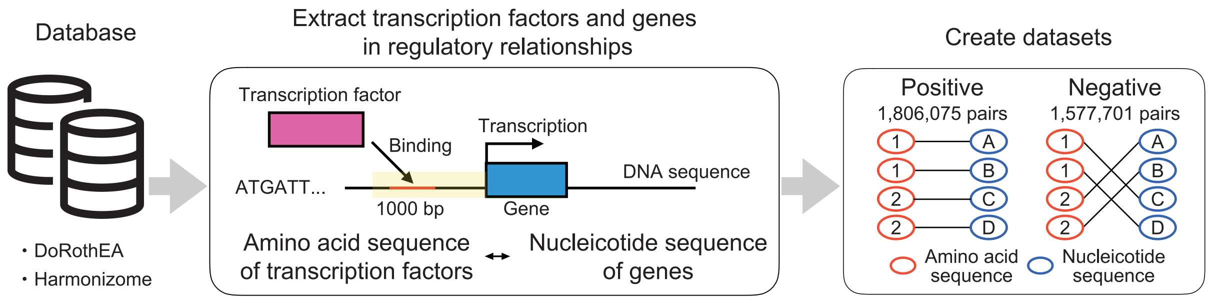


Figure S1: Overview of dataset construction

We obtained paired data for transcription factors (TFs) and target genes from the DoRothEA and Harmonizome databases. We collected the TF amino acid sequences and the target gene sequences (upstream 1000 bp of each gene) from the public genome database GenBank (accession number: GCF_000001405.39) and paired them. Cleaned paired data (see the Methods section) were labeled as positive. We then shuffled the paired data and assigned negative labels to the pairs for which no gene regulatory relationships were detected.

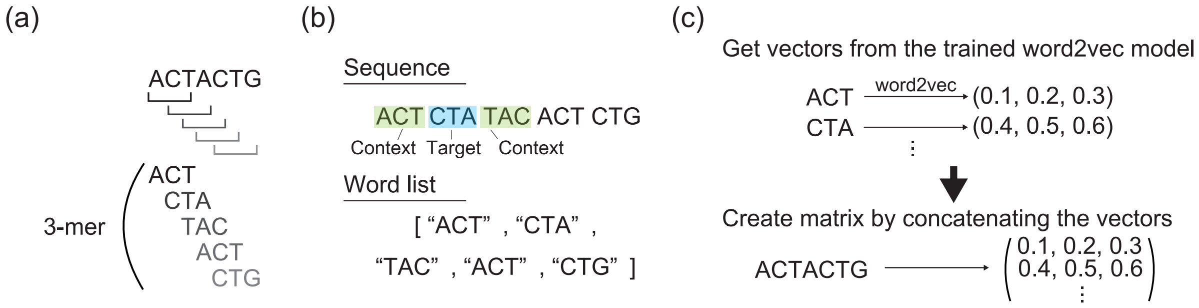


Figure S2: Procedure for obtaining an embedded matrix representation of amino acid sequences and nucleotide sequences for $k = 3$

(a) Sequences were split into $k$-mers (sub-sequences of a length k amino acids or nucleotides). (b) A word list was constructed in which all 3-mers were arranged as words in a sentence. In the word2vec model with the skip-gram algorithm, for example, " CTA " is used as the target word, and " ACT " and " TAC " are used as the context. The word2vel model was trained to predict the target word from the context. (c) Using pre-trained word2vec, we obtained a numerical vector for each 3-mer. By concatenating the numerical vectors, we obtained an embedded numerical matrix.

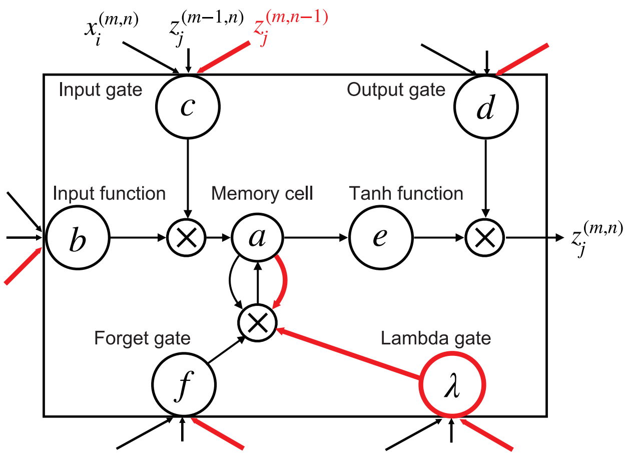


Figure S3: Architecture of 2D LSTM

Black lines, architecture common to 1D and 2D LSTM; red lines, architecture specific to 2D LSTM; $a$, a memory cell; $b$–$f$ and $\lambda$, units: $b$ receives external inputs, $c$ determines the value of the input gate, $d$ determines the value of the output gate, $e$ applies an activation function to the output of the memory cell, $f$ determines the value of the forget gate, and $\lambda$ determines the value of the lambda gate. In 2D LSTM, the output of the input layer $x_i^{(m,n)}$ at any given time step $(m, n)$ and the outputs of the memory units from one time step ago $z_j^{(m-1,n)}$, $z_j^{(m,n-1)}$ were input to each unit.

Table S1: Hyperparameters in the proposed model

To tune hyperparameters in our model, we performed Bayesian optimization using Tree-structured Parzen Estimator algorithm.

| | |
|---|---|
| Feature extraction section for amino acid sequences | |
| Kernel size in the convolution layer | 30 |
| Stride in the convolution layer | 4 |
| Number of channels in the convolution layer | 300 |
| Kernel size in the pooling layer | 15 |
| Drop-out probability after the convolution layer | 0.458280 |
| Feature extraction section for nucleotide sequences | |
| Kernel size in the convolution layer | 18 |
| Stride in the convolution layer | 4 |
| Number of channels in the convolution layer | 260 |
| Kernel size in the pooling layer | 15 |
| Drop-out probability after the pooling layer | 0.074344 |
| Feature mixing section | |
| Number of input channels to the 2D LSTM architecture | 190 |
| Number of dimensions in the intermediate layer in the 2D LSTM architecture | 190 |
| Drop-out probability after the 2D LSTM architecture | 0.136456 |
| Number of dimensions in a fully connected layer | 140 |

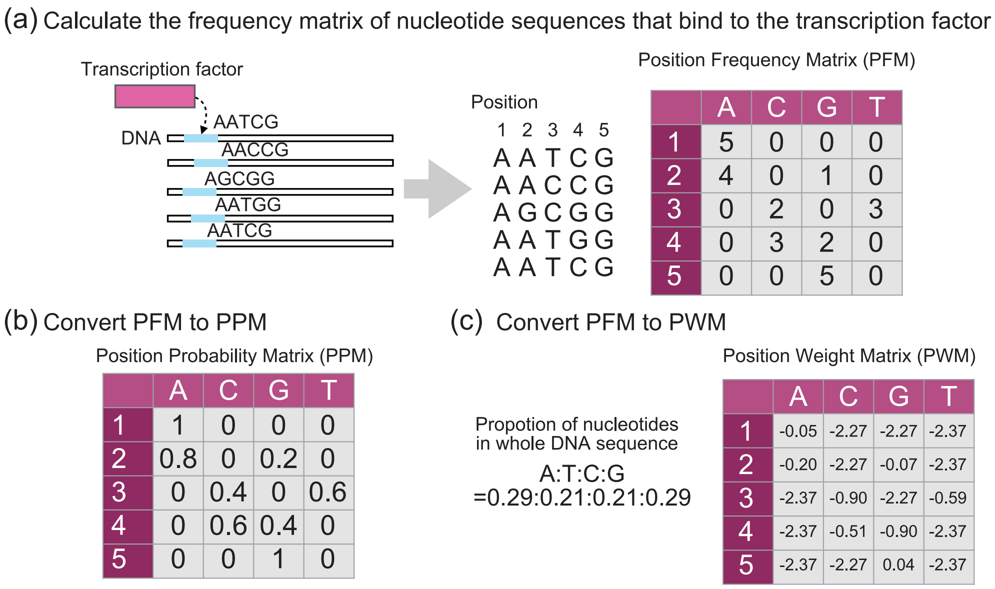


Position Frequency Matrix (PFM)

| | A | C | G | T |
|---|---|---|---|---|
| 1 | 5 | 0 | 0 | 0 |
| 2 | 4 | 0 | 1 | 0 |
| 3 | 0 | 2 | 0 | 3 |
| 4 | 0 | 3 | 2 | 0 |
| 5 | 0 | 0 | 5 | 0 |

Position Probability Matrix (PPM)

| | A | C | G | T |
|---|---|---|---|---|
| 1 | 1 | 0 | 0 | 0 |
| 2 | 0.8 | 0 | 0.2 | 0 |
| 3 | 0 | 0.4 | 0 | 0.6 |
| 4 | 0 | 0.6 | 0.4 | 0 |
| 5 | 0 | 0 | 1 | 0 |

Position Weight Matrix (PWM)

| | A | C | G | T |
|---|---|---|---|---|
| 1 | -0.05 | -2.27 | -2.27 | -2.37 |
| 2 | -0.20 | -2.27 | -0.07 | -2.37 |
| 3 | -2.37 | -0.90 | -2.27 | -0.59 |
| 4 | -2.37 | -0.51 | -0.90 | -2.37 |
| 5 | -2.37 | -2.27 | 0.04 | -2.37 |

Figure S4: PWM calculation

(a) Experimental data stored in the public DoRothEA and Harmonizome databases were used to obtain all TF-binding sites in gene nucleotide sequences. To calculate a position frequency matrix (PFM), the frequency of occurrence at each position was determined. (b) To calculate the position probability matrix (PPM), the element values of the PFM were normalized to the number of sequences. (c) The position weight matrix (PWM) was obtained by correcting the bias in the occurrence probability of the sequence in the entire gene nucleotide sequences. The equations are provided in the Methods section.

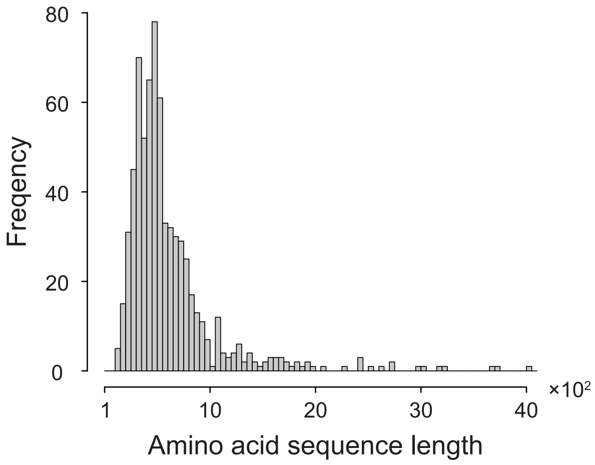


Figure S5: Sequence length distribution of TFs in the training data

Sequence length and frequency refer to TFs. The groups of sequence lengths were constructed in increments of 50 residues.

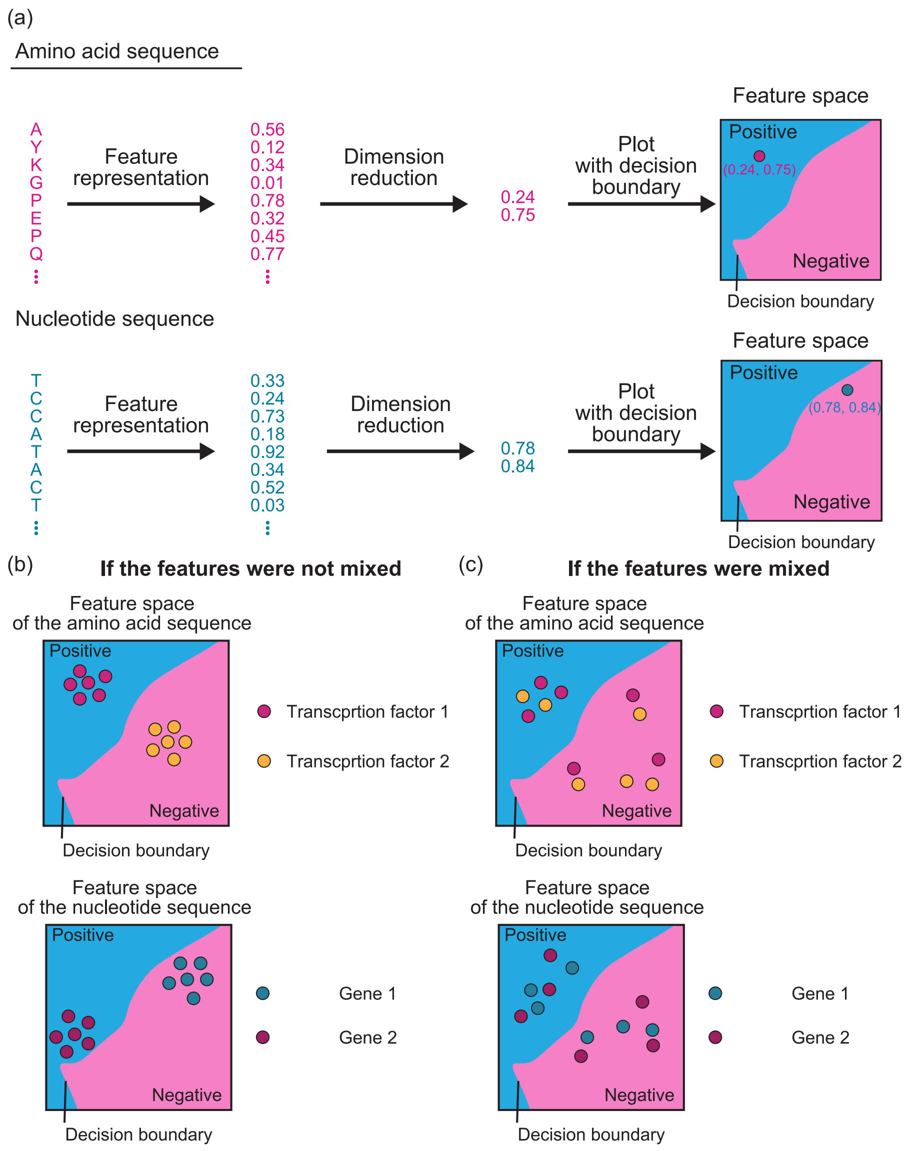


Figure S6: Conceptual diagram of the feature mixing for TF amino acid sequences and target gene nucleotide sequences

(a) Features extracted from the TFs and genes are mixed using a 2D LSTM. To qualitatively evaluate whether feature mixing contributed to the prediction or not, we performed dimension reduction for the TF and gene features, and visualized the distribution of the features. Regulatory relationships are found in blue regions but not in red regions. (b) If the features were not mixed in 2D LSTM, the features of each TF or gene would be aggregated in the same area without crossing the decision boundary for the presence or absence of a gene regulatory relationship. (c) If the features were mixed in 2D LSTM, the features of each TF or gene would be distributed across the decision boundary.

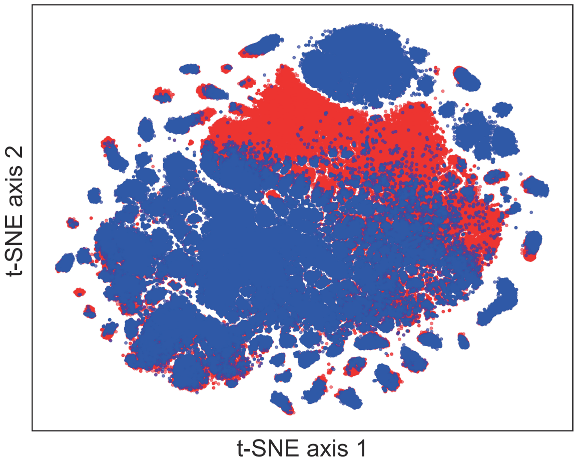


Figure S7: Dimension reduction of the features obtained from the last layer of our model using t-SNE. Each dot corresponds to each input datapoint. Blue, positive; red, negative.
